# Cone-driven, geniculo-cortical responses in canine models of outer retinal disease

**DOI:** 10.1101/2023.12.13.571523

**Authors:** Huseyin O. Taskin, Jacqueline Wivel, Gustavo D. Aguirre, William A. Beltran, Geoffrey K. Aguirre

## Abstract

**Purpose:** Canine models of inherited retinal degeneration are used for proof-of-concept of emerging gene and cell-based therapies that aim to produce functional restoration of cone-mediated vision. We examined functional MRI measures of the post-retinal response to cone-directed stimulation in wild type (WT) dogs, and in three different retinal disease models.

**Methods:** Temporal spectral modulation of a uniform field of light around a photopic background was used to target the canine L/M (hereafter “L”) and S cones and rods. Stimuli were designed to separately target the post-receptoral luminance (L+S) and chrominance (L–S) pathways, the rods, and all photoreceptors jointly (light flux). These stimuli were presented to WT, and mutant *PDE6B*-RCD1, *RPGR*-XLPRA2, and *NPHP5*-CRD2 dogs during pupillometry and fMRI.

**Results:** Pupil responses in WT dogs to light flux, L+S, and rod-directed stimuli were consistent with responses being driven by cone signals alone. For WT animals, both luminance and chromatic (L–S) stimuli evoked fMRI responses in the lateral geniculate nucleus (LGN) or visual cortex; RCD1 animals with predominant rod loss had similar responses. Responses to cone-directed stimulation were reduced in XLPRA2 and absent in CRD2. NPHP5 gene augmentation restored the cortical response to luminance stimulation in a CRD2 animal.

**Conclusions:** Cone-directed stimulation during fMRI can be used to measure the integrity of luminance and chrominance responses in the dog visual system. The *NPHP5*-CRD2 model is appealing for studies of recovered cone function.

**Translational Relevance:** fMRI assessment of cone driven cortical response provides a tool to translate cell/gene therapies for vision restoration.

## Introduction

The dog is an important model for studies of retinal cone disease and its treatment, owing to the availability of diverse, naturally occurring genetic disorders and to the presence of a fovea-like, cone-rich zone in the canine area centralis^1^. The ultimate goal of treatment—whether by somatic-cell gene therapy^2,3,4,5^ or transplantation of cone precursors^6,7^—is restoration of function, with vision supported by the cones under day-light conditions of particular importance. Measurement of visual behavior in post-treatment animals, however, requires training that can be difficult to start prior to successful therapy. Further, measuring the effect of treatment is complicated in some models by the presence of residual cone function at baseline.

Non-invasive measurements of physiologic and neural response have been used as a proxy for visual function. Flash electroretinography (ERG) provides a measure that principally reflects responses of the photoreceptors and middle retina, integrated across eccentricity. Similarly, pupil constriction to light probes spatially integrated retinal responses and the brainstem reflex circuit. While methodological refinements can provide spatial maps of retinal function and target cell classes, it remains the case that ERG and pupillometry are unable to confirm that restored retinal function drives signals in the cortical visual pathway. This is of particular importance for studies of cell therapy, as the ability of transplanted photoreceptor precursor cells to re-create functional connections with the inner retina is still to be determined^6^.

Visual evoked potentials (VEP) measure the cortical response to visual stimulation and have been used as a measure of treatment response in canine retinal gene therapy^8^. VEPs are challenging to measure in the dog, however, due to the small signal, risk of contamination from ERG responses, and the sensitivity of the measurement to anesthesia^9^. Cortical function may also be measured in the dog using functional magnetic resonance imaging (fMRI)^10^. We have previously shown that retinal gene therapy in *RPE65*-LCA is associated with a restoration of fMRI responses from the canine visual cortex^11^. In this prior study, visual stimulation was a high-photopic, flashing screen that alternated with periods of darkness. Consequently, both rod and cone responses could have contributed to the measured cortical activity.

In the current study we measured with fMRI cortical responses to cone-directed stimulation. In dichromatic mammals like dogs, signals from the cones contribute to two post-receptoral pathways that encode luminance (overall brightness) and chrominance (“blue-yellow”) (Figure 1a). One goal of the current study was to determine if separate responses from the luminance and chromatic pathways are measurable in dogs with no retinal disease. We also collected data from animals with each of three different inherited retinal diseases that differ in the extent of rod or cone involvement. Our results demonstrate that fMRI provides a reliable, within-animal assessment of the presence of post-retinal cone signals in the dog, and that a particular disease model (*NPHP5*-CRD2) is especially promising for the study of recovered cone function.

**Figure 1:**
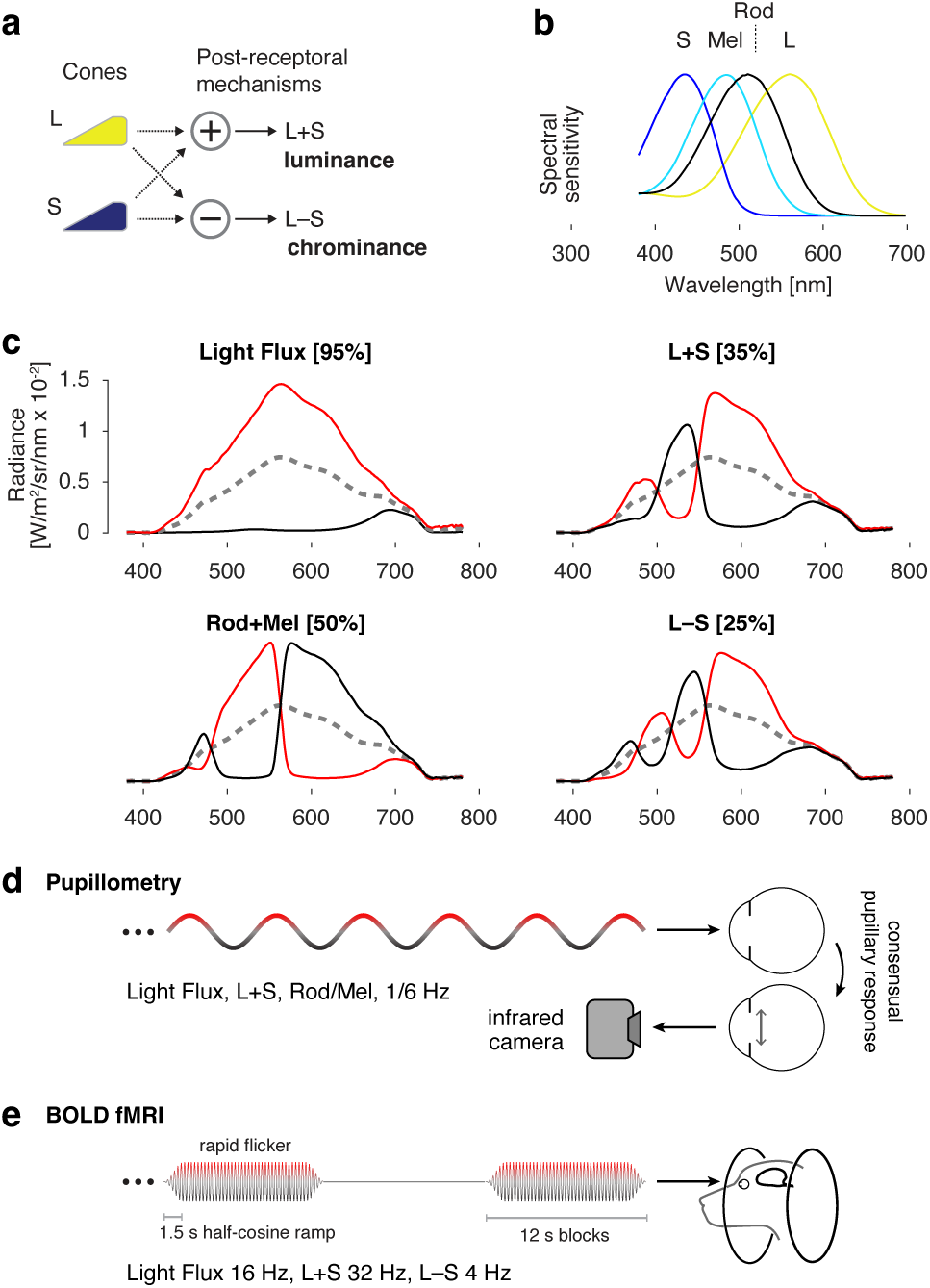
a) Signals from the L/M (hereafter “L”) and S cones are combined and contrasted to create two post-retinal channels that signal luminance and chrominance. b) Spectral sensitivities of the canine photoreceptors. c) Stimulus spectra used to target particular photoreceptor combinations. Each stimulus modulated between a stimulating (red) and suppressing (black) spectrum around a common background (dashed gray). The nominal contrast on the targeted photoreceptors is given in brackets. Light flux produces equal contrast on all photoreceptors. d) A spatially uniform field of light was presented to one eye. The field modulated between a stimulating and a suppressing spectrum, following a sinusoidal temporal waveform. In measurements of pupil response the modulation frequency was 1/6 Hz, and the consensual pupil response to the stimulus was recorded from the unstimulated eye using an infrared camera. e) In fMRI measurements a higher frequency flicker was presented in 12 second blocks, interleaved with presentations of the static, photopic background spectrum.

## Methods

### Subjects

The animals evaluated were part of a canine research colony maintained at the University of Pennsylvania, Retinal Diseases Studies Facility. All procedures were carried out in strict accordance with the ARVO Statement for the Use of Animals in Ophthalmic and Vision Research and approved by the Institutional Animal Care and Use Committee of the University of Pennsylvania (IACUC no. 803254). The study examined WT dogs, and dogs affected with late-stage inherited retinal degeneration caused by mutations in *PDE6B* (RCD1 model)^12^, *RPGR* (XLPRA2 model)^13^, and *NPHP5/IQCB1* (CRD2 model)^14^. None of the dogs in this study were neutered. A description of the studied animals is provided in Table 1, and the known impairments in retinal and visual function in these disease models are described in Supplemental Table S1. To summarize: RCD1 has severe rod loss with relatively preserved cone function, while XLPRA2 and CRD2 both have diminished cone function, with this loss being nearly complete in CRD2.

**Table 1:** Summary of the studied animals.

| Animal ID | Sex | Disease Type | Age at pupillometry | Age at fMRI | Pupillometry doses (max mg/kg/hr) |  |  | fMRI doses (max mg/kg/hr) |  |  |
| --- | --- | --- | --- | --- | --- | --- | --- | --- | --- | --- |
|  |  |  |  |  | Ketamine | Midazolam | Dexmedetomidine | Ketamine | Midazolam | Dexmedetomidine |
| N344 | M | WT | 8 months | 14 months | 15 | 0.6 | 0.002 | 14 | 0.6 | 0.0028 |
| N349 | M | WT | 12 months | 21 months | 14 | 0.6 | 0.0016 | 13 | 0.5 | 0.00175 |
| N347 | F | WT | 6 months* | 23 months | 52 | 0.6 | 0.03 | 13 | 0.5 | 0.00225 |
| M662 | F | WT | - | 120 months | - | - | - | 15 | 0.6 | 0.003 |
| 2350 | M | RCD1 | 8 months | 7 months | 9 | 0.4 | 0.003 | 12 | 0.5 | 0.003 |
| 2353 | M | RCD1 | 6 months | 15 months | 8 | 0.6 | 0.002 | 15 | 0.6 | 0.003 |
| 2356 | F | RCD1 | 8 months | 9 months | 9 | 0.4 | 0.003 | 14 | 0.6 | 0.002 |
| 2346 | F | RCD1 | - | 28 months | - | - | - | 11 | 0.6 | 0.00175 |
| Z663 | M | XLPRA2 | 6 months | 9 months | 13 | 0.5 | - | 13 | 0.6 | 0.003 |
| Z665 | M | XLPRA2 | 8 months | 9 months | 15 | 0.6 | 0.002 | 10 | 0.5 | 0.003 |
| Z666 | F | XLPRA2 | 10 months | 9 months | 11 | 0.5 | 0.003 | 10 | 0.5 | 0.001 |
| Z709 | M | XLPRA2 | - | 18 months | - | - | - | 12 | 0.6 | 0.002 |
| Z710 | M | XLPRA2 | - | 18 months | - | - | - | 10 | 0.6 | 0.002 |
| AS2-454 | F | CRD2 | - | 5 months | - | - | - | 10 | 0.5 | 0.002 |
| AS2-451 | M | CRD2 | - | 5 months | - | - | - | 10 | 0.5 | 0.002 |
| WM65 | M | CRD2 | - | 7 months | - | - | - | 10 | 0.5 | 0.002 |
| WM67 | F | CRD2 post tx | - | 10 months | - | - | - | 10 | 0.6 | 0.0025 |

\* Suitable anesthesia levels could not be achieved for this animal during pupillometry

The primary study collected fMRI data from 3 RCD1, 5 XLPRA2, 3 CRD2, and 3 WT animals; data from a fourth RCD1 animal (ID:2353) was collected but was unusable due to a temporal noise source of unclear cause. A subset of these animals was also studied with pupillometry: 3 RCD1, 3 XLPRA2, and 2 WT (one WT animal could not be studied due to the inability to achieve a suitable level of anesthesia).

An additional CRD2 animal was studied with fMRI 32 weeks following retinal gene therapy to one eye^5^. This same animal had undergone ERG and visual behavior assessment in an obstacle avoidance course 24 weeks post-treatment using previously described methods^15^. Finally, an additional WT animal was studied with a different, “localizer” fMRI protocol for the purposes of defining the anatomical location of the lateral geniculate and visual cortex.

### Anesthesia and Animal Preparation

The animals were given subcutaneous acepromazine (0.25 mg/kg) and atropine (0.02 mg/kg) prior to transportation to the experimental site. The time between the administration of this drug and induction varied, but was at least 1.5 hours. Anti-nausea medication (maropitant citrate, Cerenia®, Zoetis) was given 20 minutes prior to induction. Induction was performed with ketamine (1-10 mg/kg IV) and midazolam (0.2-0.5 mg/kg IV). A loading dose of dexmedetomidine (2-5 μg/kg) was used, and anesthesia was maintained during the experiment with ketamine (5-15 mg/kg/hr IV), midazolam (0.2-0.6 mg/kg/hr IV), and dexmedetomidine (1-3 μg/kg/hr IV). The functional MRI studies were conducted under pharmacologic paralysis; vecuronium was given as a loading (0.1 mg/kg) and then a maintenance dose (0.1-0.2 mg/kg/hr) during scanning. Paralysis was not used for the pupillometry studies.

During all data collection, animals were intubated and mechanically ventilated with 100% O_2_. Respiratory rates that were multiples of 5 breaths per minute were avoided to avoid the introduction of cardiopulmonary signals at harmonics of the fMRI stimulus frequency. Pulse oximetry, end-tidal CO_2_, and core body temperature were monitored throughout data collection.

For the functional MRI studies, both eyes were pharmacologically dilated with topical atropine, tropicamide, and phenylephrine once when the dogs arrived at the experimental site and again 20-30 minutes before induction; pupil dilation was not performed for the pupillometry studies as the goal was to measure the consensual pupil response. The eyelids were held open during fMRI studies with sprung plastic specula, and during pupillometry with lid sutures. The eyes were lubricated at frequent intervals throughout the study with artificial tears. Earplugs and foam padding over the ears were provided during fMRI scanning. Room lights were turned off.

### Spectral modulations

Light stimuli were produced by a digital light synthesis engine (OneLight Spectra), conveyed via a fiber optic cable, and then presented to the animal within a custom-made, MRI-compatible, circular eyepiece with an ∼26-degree uniform visual field^16^. Intermittent calibration of the device and regular measurements of the spectral properties of the stimuli was performed with a spectroradiometer (PhotoResearch PR-670). During data collection, the eyepiece was held with an articulated plastic arm and positioned ∼5 mm from the corneal surface of the stimulated eye. The fellow eye was uncovered during data collection. During the pupillometry sessions, an infrared video camera (640×480 pixel resolution at 60 Hz interlaced; LiveTrack, Cambridge Research Systems) was used to record the pupil response from the non-stimulated eye.

The stimuli for both experiments targeted particular photoreceptor classes using the method of silent substitution^17,18^. Stimuli were designed to stimulate or silence the L cone (also termed the “L/M” or “ML” cone in dichromatic animals), S cone, rod, and melanopsin photopigments (Figure 1b). The spectral sensitivity of these photoreceptor classes was modeled using the Govardovskii nomogram^19^, with a lambda max of 555, 429, 506, and 480 nm, for the L, S, rhodopsin, and melanopsin photopigments respectively. The spectral transmittance of the canine crystalline lens was included in the calculation to account for pre-receptoral filtering^20^. A non-linear search across device settings was used to construct spectral modulations that had the desired property of silencing some photoreceptors while maximizing contrast on targeted photoreceptors (Figure 1c). The modulations were presented around a common, half-on spectral background. Neutral density filters were placed in the light path to bring the background into the desired luminance range. The mean (±SD) human luminance of the stimulus across experiments was 438 ± 83 cd/m^2^ for pupillometry, and 305 ± 92 cd/m^2^ for MRI, which corresponds to corneal irradiance of 0.29 ± 0.05 and 0.20 ± 0.06 Watts/m^2^ respectively. Spectroradiometric measurements of the stimuli were made immediately before and after each data collection session. Table S2 provides the calculated contrast on the targeted and nominally silenced photoreceptors.

### Pupillometry stimulus and data analysis

In each of many acquisitions, a bipolar modulation of the spectral content of the stimulus was presented to one eye, while the consensual pupil response was recorded from the fellow eye under closed-loop conditions. The modulation followed a slow (1/6 Hz) sinusoidal temporal profile of transition from one arm of the spectral pair to the other, passing through the background spectrum (Figure 1d). Each acquisition was 360 seconds in duration. Three acquisitions were obtained in order using the L+S, Rod+Mel, and Light Flux modulations. The stimulated and recorded eyes were then switched, and this set of acquisitions was repeated. A total of 12 acquisitions were made in a measurement session for a given animal. An external transistor-transistor logic (TTL) pulse was used to initiate both the stimulus and the video recording, establishing a common temporal reference.

The resulting IR videos were processed using previously described open-source software^21,22^. The primary operation of the analysis was to fit an ellipse to the contrast-defined border between the pupil and iris for each frame of the video. Imperfections in the border segmentation (due to, e.g., the presence of eyelashes or the first Purkinje reflex of the active IR source) were removed by iteratively constraining the parameters of the ellipse and removing poorly-fit border points. In some cases, translational motion correction was applied to the images to correct head motion during the recording interval. Acquisitions in which >50% of video frames contained poor pupil fits were excluded from the analysis; 4 acquisitions were so excluded, out of the total of 96 acquisitions collected.

The ellipse area over time was expressed as percentage change and the time series data were fit by linear regression with a sine and cosine at 1/6 Hz. The resulting amplitude and phase of the fit were obtained. The standard error of the mean of these fits was estimated by repeating the fit across an exhaustive bootstrap resampling of all 35 available combinations of the four measurements sampled with replacement and taking the standard deviation of the set of values.

### MRI stimulus and image pre-processing

In each of many functional MRI acquisitions, a rapidly flickering spectral modulation was presented to one eye of the animal. The flicker was presented in 12-second blocks, alternating with 12 seconds of the steady, photopic background spectrum. The flicker within each stimulation block was subject to a 1.5-second half-cosine temporal window at onset and offset (Figure 1e). Each acquisition was 432 seconds in duration. Three acquisitions were obtained, in order, using the L+S, L–S, and light flux modulations. The flicker frequency of these stimuli was 32 Hz, 4 Hz, and 16 Hz, respectively, selected based upon the temporal sensitivity of canine vision^23,24^. The set of three acquisitions was collected for one eye, then the stimulus was switched to the other eye, and the acquisition set was repeated. A total of 18 acquisitions were acquired in a given scanning session.

MR imaging was performed on a 3T Trio Siemens scanner with a 15-channel knee coil. Two MPRAGE images were collected at the beginning of each scan with the *tfl3d1 sequence, 0.729 x 0.7 x 0.729 voxel size, TR=1700ms, TE=4.87ms, and flip angle=9°. The BOLD images were collected with the epfid2d1_64 sequence, 2×2×3 voxel size, TR=3000ms, TE=30ms, and flip angle=90°. Immediately after acquiring the MPRAGE images, two single-TR scout images with the same parameters as the BOLD images were obtained with alternating AP and PA phase encoding directions, and these were used for susceptibility distortion correction. The main BOLD sequences were also collected with alternating phase encoding directions.

For each subject, the two structural images were bias-corrected with N4 Bias Correction^25^ and spatially aligned with linear registration. The registered images were averaged. An initial approximate brain mask was created on the resulting image with FSL Brain Extraction Tool^26^. This mask was used as an input to the altAntsBrainExtraction algorithm to perform skull stripping (https://github.com/cookpa/altAntsBrainExtraction). The skull-stripped image was then warped to a canine template^27^ with non-linear diffeomorphic (SyN) registration^28^.

A susceptibility distortion field map was calculated from the AP and PA scout image pairs using FSL top-up^29^. This field map was used to perform distortion correction on the scout and fMRI time-series images. The resulting time series images were motion corrected with rigid-body registration using the middle TR volume as the target^30^, and the mean volumetric image across time was obtained from the motion-corrected data. This mean volume was linearly registered to one of the top-up corrected scout images, and then warped to the canine template with a single interpolation using the warp fields that were previously calculated between the structural and template coordinates.

Occasional transient, high-amplitude “spikes” were observed in the raw time series data in some sessions (attributed to electrical arcing at the coil plug sockets). We created an algorithm to detect these spikes. First, the linear trend was removed from the raw functional images, and then outlier detection was performed for each voxel across time by calculating the median absolute deviation (MAD). A voxel was considered an outlier if it was at least 6 MAD above the median. This detection resulted in a vector containing the number of noisy voxels for each TR in an image. With another outlier detection performed on this vector, we identified time points whose number of noisy voxels was at least 25 MAD above the median value and created confound regressors for these time points.

Although the animal was paralyzed during scanning, brain motion was still present due to cardiopulmonary activity. For each acquisition, twenty-four motion regressors were obtained from the least-squares motion correction operation (original parameters, temporal derivatives, and squares of both). A principal components analysis was then performed upon this set of regressors and a reduced set of vectors that explained 95% of the variance was obtained. Additional vectors for each image spike were added, and the variance attributable to the resulting confound matrix was regressed from each voxel.

As we used a uniform-field, monocular stimulus, and given that the dog has approximately 60° of binocular vision^31^, we might expect that post-chiasmal neural responses would be larger in the eye contralateral to the stimulus as compared to ipsilateral. Consistent with this, we found a reliable difference in cortical response between the two hemispheres in WT animals in response to the light flux stimulus (Figure S1). While the overall magnitude of response may differ between the hemispheres, we had no *a priori* reason to think that this difference would interact with the effect of photoreceptor direction or disease model. Therefore, we combined responses from the left and right brain hemispheres by mirror-reversing the preprocessed time series data around the sagittal plane, performing a linear registration between the original and flipped images, and then averaging them together.

### fMRI Model Fitting

The fMRI time-series data from each imaging session was fitted using an open-source, non-linear model fitting routine (https://github.com/gkaguirrelab/forwardModel). For each session, the set of 6 acquisitions that were obtained for a particular stimulus type were concatenated, combining across left and right eye stimulation. The model consisted of a square-wave representation of the 12-second on and off blocks of the stimulus, with the amplitude of this response under the control of a different parameter for each of the 6 acquisitions. The square wave representation was then convolved with a model of the hemodynamic response function, itself defined by three parameters that describe a weighted combination of a three-component, hemodynamic basis set^32^. A non-linear search was performed to identify model parameters that minimized the L2 norm of this model fit to the observed data in each voxel. The mean amplitude of response (beta) was obtained and averaged for each disease model and stimulus modulation. An additional analysis was performed for data averaged across voxels within pre-defined ROIs (defined below) and results were plotted for each subject separately.

### Region of interest definition

We defined visually-responsive regions of interest using localizer data collected from a separate WT animal. The stimulation used for this localizer measurement was the 12-second, blocked 16 Hz light flux flicker around a photopic (∼400 cd/m^2^) background, alternating with 12-second periods of darkness (∼1 cd/m^2^). A total of 44 acquisitions were obtained in 3 separate sessions. Anesthesia, MR imaging parameters, data preprocessing, and statistical analysis were performed as described above and a map of the R^2^ of the model fit to the across-acquisition, average time-series was obtained. The resulting map was thresholded and binarized, yielding two discrete regions corresponding to the visual cortex (143 voxels in size), and the lateral geniculate nucleus (27 voxels in size). These two regions were then used to interrogate the data collected from other studied animals.

## Results

### Pupil responses to cone-directed stimuli show minimal signs of rod intrusion

We measured consensual pupil responses to slow (1/6 Hz) sinusoidal modulations of the spectral content of a field of light presented to one eye. The first of these modulations, “light flux”, produces high-contrast, joint stimulation of the cones, rods, and melanopsin, although the photopic stimulus background might be expected to saturate rod responses, and the stimulus frequency is fast relative to melanopsin kinetics^16^. A clear pupillary response at the stimulation frequency was seen in the across-animal, average response for each of the three studied groups (Figure 2, top row).

**Figure 2:**
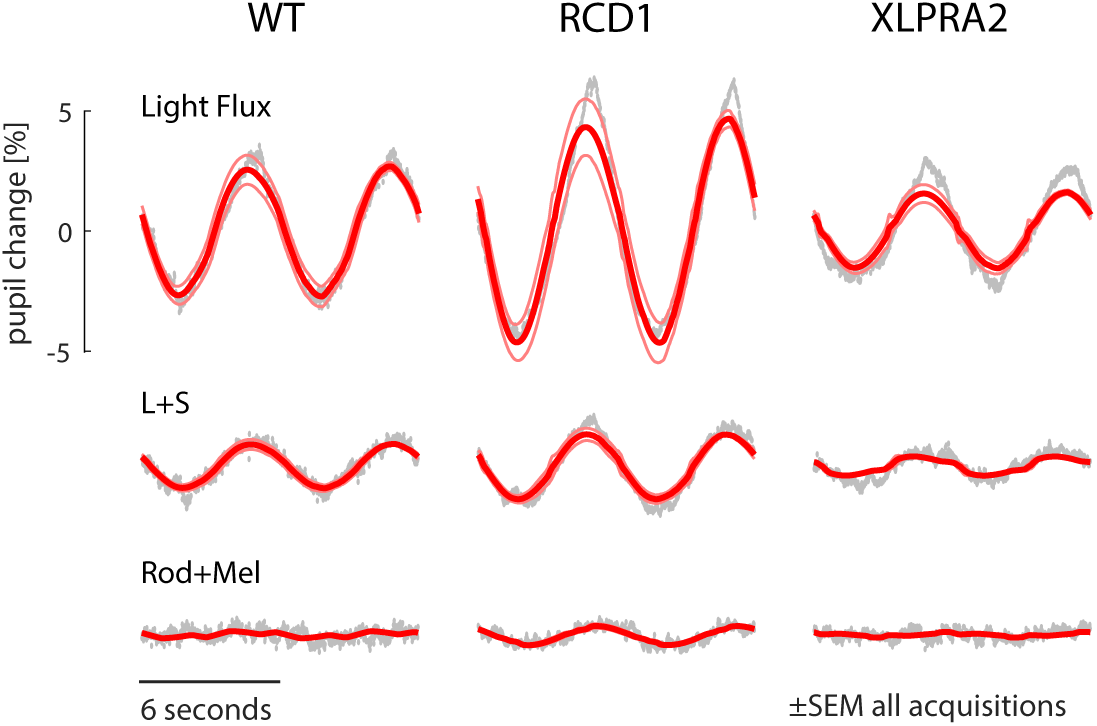
Mean (across animals) pupil response to three different stimulus conditions: light flux (which drives all photoreceptor classes), L+S (cone luminance, silencing the rods and melanopsin), and Rod+Mel (silencing the cones). Measurements were made for three groups of animals (WT, RCD1, and XLPRA2). The measured pupil response is in gray, and the fit of a sinusoidal response model is in red, with the thin lines indicating the standard error of the mean across all acquisitions from all animals in a group.

Next, we measured the pupil response to a modulation designed to produce isolated L and S cone stimulation. Because of the need to tailor the spectral modulation to nominally silence the rods and melanopsin, the contrast of this stimulus upon the cones was about a third of that produced by light flux (35% vs 95%). This L+S stimulus also evoked a pupillary response in all three disease models, although relatively reduced in XLPRA2 (Figure 2, middle row). As would be predicted by the decrease in stimulus contrast, the amplitude of the pupil response to the L+S stimulus was roughly 30% of that produced by the light flux stimulus (ratio of L+S response amplitude to light flux response amplitude: 0.34 in WT; 0.28 in RCD1; 0.25 in XLPRA2).

Finally, we measured the response to a stimulus that produced substantial (50%) contrast on the rods and melanopsin while silencing the cones. There was effectively no response in the WT and XLPRA2 groups, and only a small response was observed in RCD1.

Figure 3 presents the measured response for each animal and stimulus condition in a polar plot representation. While there was considerable variation between individual animals in the amplitude of response, phase was consistent across animals for the light flux and L+S stimulus conditions. No difference was found in phase between the light flux and L+S stimuli across animals (paired t-test [7 df] = 0.21, p = 0.84). This suggests that slower melanopsin and rod signals make a minimal contribution to the light flux response^16,33^. Responses to the Rod+Mel condition were of low amplitude and variable phase.

**Figure 3:**
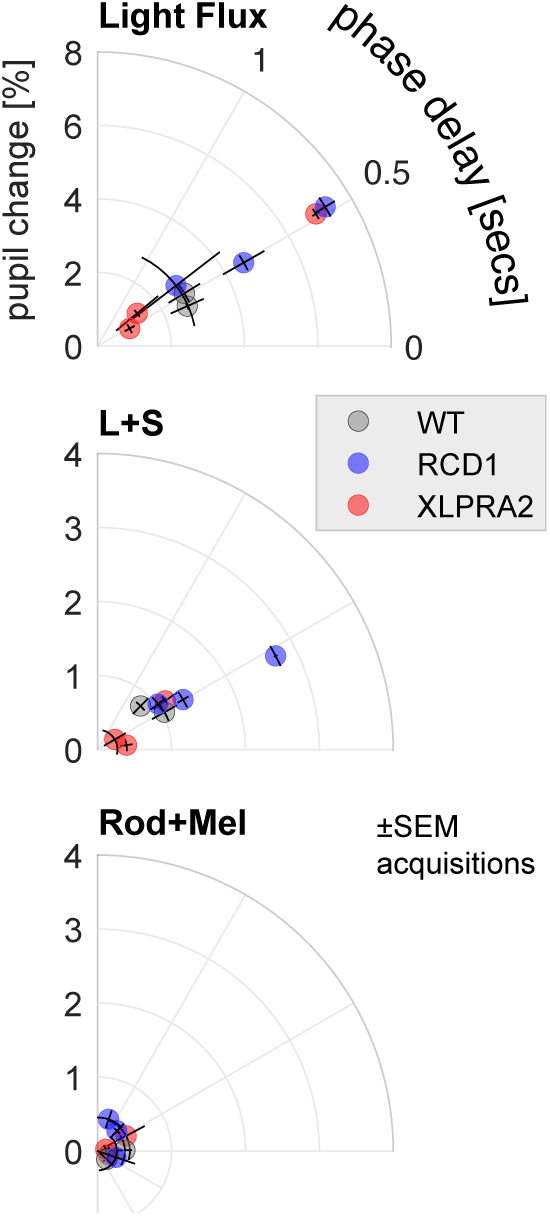
Polar representation of pupil responses to the three stimulus conditions for each studied animal. Distance from the origin indicates the amplitude of pupil change, and angular position indicates the phase delay. The SEM of the response measured for each animal (across acquisitions) is indicated by the error bars.

### Brain response to cone-directed stimulation differs across disease models

We created a flat-map representation of the canine posterior cortex (Figure 4) with the location of visual area borders defined by homology with the cat^34^.

**Figure 4:**
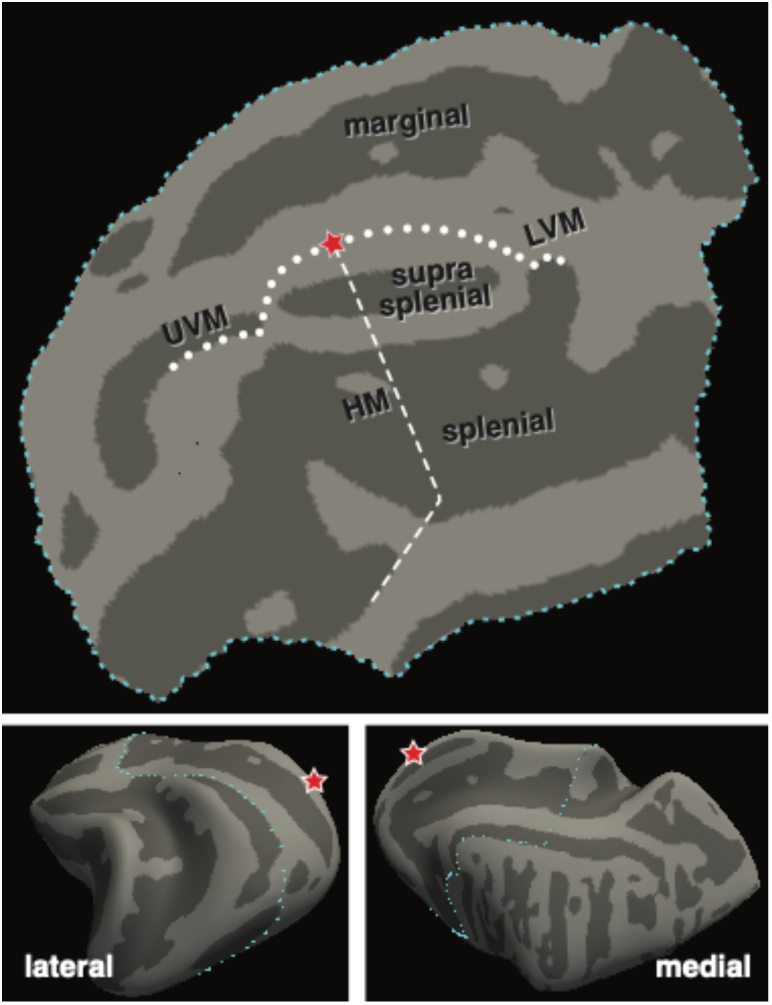
Flattened representation of the canine visual cortex. The marginal, suprasplenial, and splenial sulci are indicated. The dots show the expected position of the upper and lower vertical meridians (UVM, LVM) and the dashes show the horizontal meridian (HM). The foveal confluence is indicated with a star.

Figure 5 presents the mean fMRI response for the stimuli and studied populations. The response from the visual cortex and the lateral geniculate nucleus is shown; minimal responses were found at other anatomical sites (see Figure S2).

**Figure 5:**
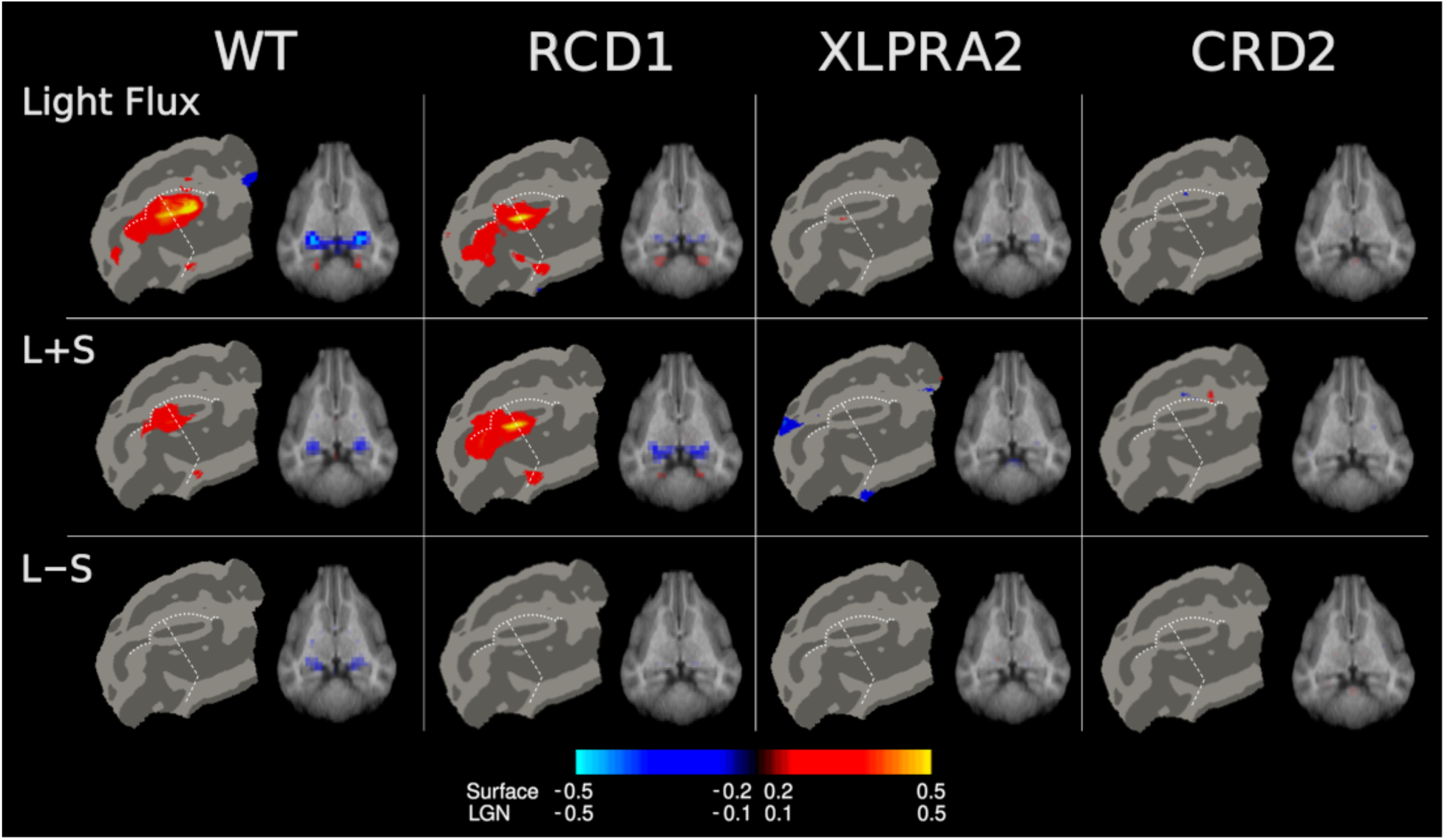
Functional MRI responses to cone-directed stimulation. The average, across animal (and hemisphere) response (percent signal change) is shown for each of three stimuli and each of four studied populations. Cortical responses are shown on a flat map, and sub-cortical response on an axial slice that passes through the lateral geniculate nucleus. Data from the two hemispheres were combined, resulting in symmetric maps. The location of the horizontal (dashes) and vertical (dots) meridians are indicated. Different map thresholds were used for the cortical surface and subcortical maps (see color bar). Responses in the cortex and thalamus are of opposite sign, perhaps due to the effects of anesthesia.

In the cortex, luminance directed stimuli (light flux and L+S) evoked fMRI responses in the vicinity of the supra-splenial sulcus, straddling the horizontal meridian and close to the vertical meridian border. This response was seen in both the WT and the RCD1 animals, both of which have intact (WT) and retained (RCD1) cone function at the studied ages. The two disease models with cone impairment (XLPRA2 and CRD2) did not have responses at this threshold. A corresponding pattern of responses in the LGN was also seen.

The isoluminant L–S stimulus did not produce a cortical response in the WT or disease model animals. There was, however, a measurable LGN response to this stimulus in the WT animal, demonstrating a response within the canine geniculo-cortical pathway to chromatic stimulation.

We quantified these respones within cortical and LGN regions-of-interest. Figure 6 presents the fMRI response for each studied animal, grouped by stimulus and disease model.

**Figure 6:**
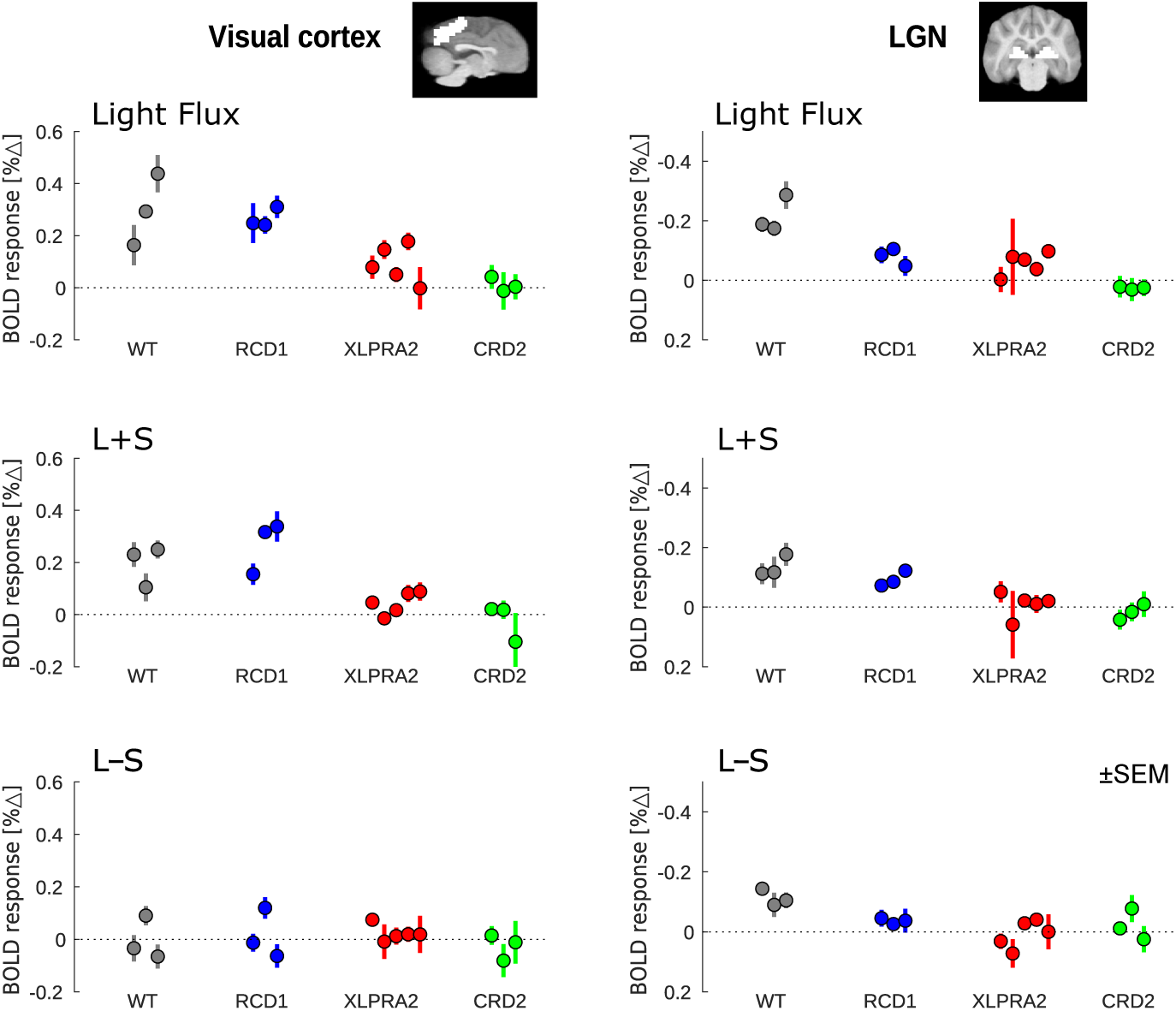
Functional MRI responses (% signal change) within the visual cortex (left) and LGN (right) for each of the three stimulus types (rows), organized by disease model. Error bars are the SEM across fMRI acquisitions obtained for an individual animal. Note the different y-axis direction and range for the cortex and LGN plots.

The light flux and L+S stimuli evoke some small responses in the XLPRA2 animals despite their extensive cone loss. In comparison, there is consistently no response to the cone-directed stimulation in the CRD2 population.

### Gene therapy restores cone-selective functional responses in the visual cortex

We studied one CRD2 animal ∼32 weeks following retinal gene therapy to the left eye. This animal received a subretinal injection at 14 weeks of age (mid-stage disease) of an AAV2/5 vector carrying a single stranded construct composed of the human *NPHP5* cDNA with a woodchuck hepatitis virus posttranscriptional regulatory element (WPRE) that was under control of the photoreceptor-specific GRK1 promoter. Figure 7 shows the cortical response in this animal to light flux stimulation around a photopic background delivered to the left eye, and to the untreated right eye. A clear difference in evoked cortical response is seen, confirming that we are able to measure recovered cone-specific function in this disease model (mean response within the cortical ROI ± SEM across acquisitions: left eye: 0.229% ± 0.094; right eye: 0.022% ± 0.086). The L+S and L–S stimuli did not evoke a reliable response in the cortex from either eye and there was no measurable response in the LGN to any of the modulation types. Full-field ERG and testing of visually-guided behavior 24 weeks after subretinal injection of the AAV2/5-*NPHP5* vector had shown in the left (treated) eye rescue of both rod- and cone-mediated function (Figs. S3, S4).

**Figure 7:**
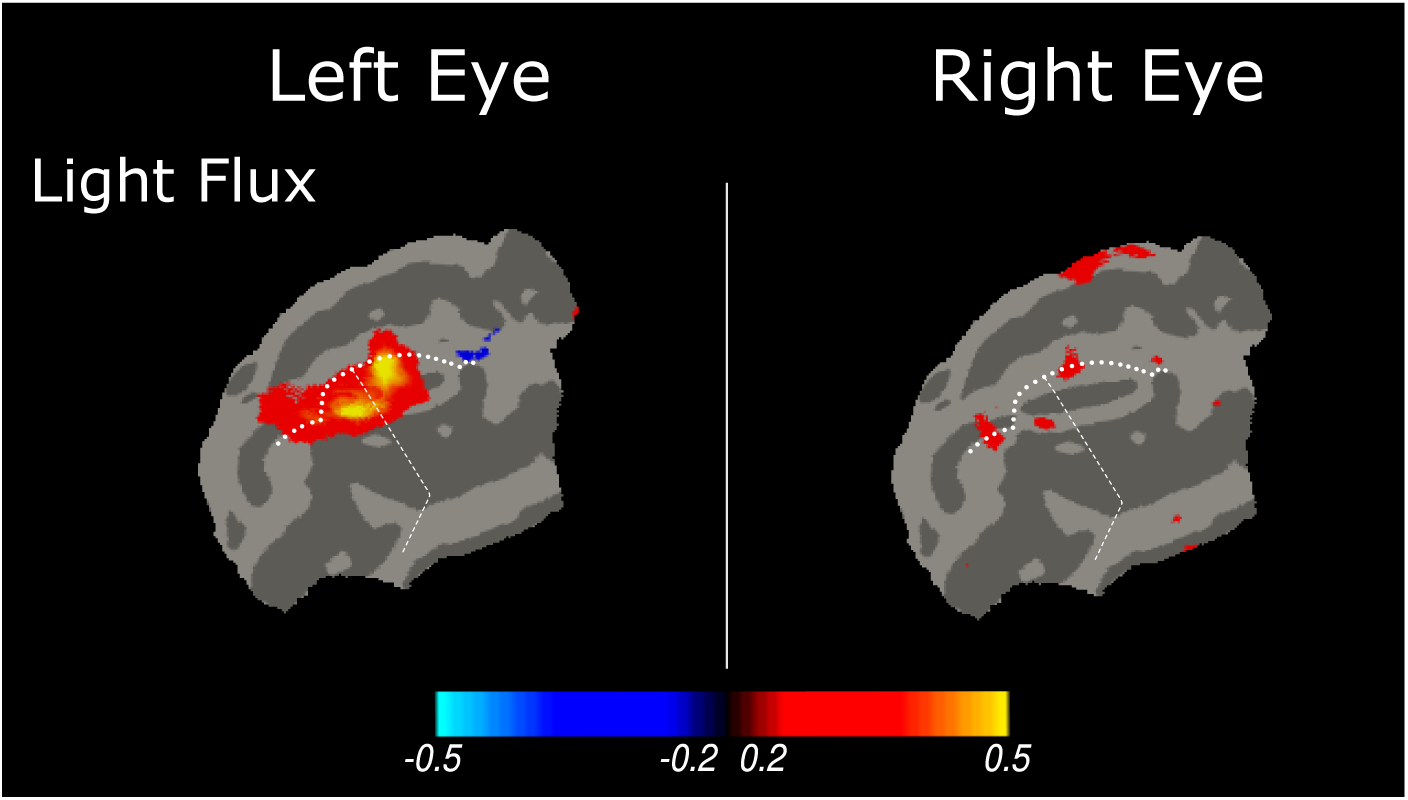
fMRI responses (percent signal change) to light flux stimulation within the visual cortex of a CRD2 animal following gene therapy delivered to the left eye.

## Discussion

Manipulation of the spectral content of stimuli has been used previously in the dog to probe photoreceptor contributions in both ERG and pupil responses^35,36^. Our work extends these techniques to fMRI measurements in normal and retinal disease models (as has been pursued in the human^37,38^). In WT dogs we find reliable responses in the geniculo-cortical pathway to both luminance and chromatic stimuli. Measurements from inherited retinal disease models support both the selectivity of our stimuli for eliciting cone-driven responses and help identify a promising disease model for the study of cone function restoration.

The three disease models we studied differ in the degree of impairment of rod and cone function. In dogs with *PDE6B*-RCD1, ∼70% of rods are lost in the outer nuclear layer by 3 months of age, but cones are relatively spared^39,40^. By the age of 6-8 months (at which we made the bulk of our measurements) further rod loss would be expected. *RPGR*-XLPRA2 affects both cones and rods and results in the loss of one-third of these photoreceptors by 3-4 months of age^4,41^, with further loss of both classes expected by the age of 6-18 months we studied. CRD2 due to *NPHP5* mutation features severe, early cone dysfunction followed by a secondary rod loss^5,42^. At the ages studied here (5-7 months) almost complete loss of rod and cone function is found.

### Normative canine visual cortex responses

The retinotopic organization of the visual cortex has not been studied in the dog, although electrophysiologic^34^ and fMRI measurements^44^ have been made in the cat. The location of maximal cortical response in our data corresponds to primary visual cortex, nearly centered on the foveal representation. While electrophysiologic data from the cat locate the fovea on the marginal gyrus, our data (and the cat fMRI study^44^) find the peak response in the adjacent supra-splenial sulcus. The elliptical extent of the cortical response we observe corresponds to ∼10° radial eccentricity^34^, which is in rough agreement with the 13° radial extent of our stimulus^45^.

We found reliable fMRI responses to chromatic (L–S) stimuli in the LGN of WT dogs, although there was not an accompanying cortical response. While the S-cones are relatively sparse in the canine retina^46^, dogs are known to be able to distinguish luminance and “blue-yellow” chrominance^47^. Our study used a relatively slow, 4 Hz L–S flicker stimulus, based upon the low-pass characteristic of the S-cone ERG response^24^ and by analogy to the temporal sensitivity of human vision for isolated S-cone modulations^48^. Visual cortex responses may have been reduced by habituation to this slow modulation. Future studies of canine chromatic responses might examine more rapid flicker, and make use of a shifted spectral background to achieve higher contrast on this channel.

### Photoreceptor targeting

We used temporal variation in the entire wavelength spectrum of our stimulus to target and nominally silence specific photoreceptor classes. The design of the stimuli, and the accuracy of photoreceptor targeting, is based upon physical measurements of our display apparatus and prior measurements of the spectral properties of the dog photoreceptor opsins and lens transmittance. We might consider the consequence of errors in stimulus targeting. Perhaps of greatest concern are false positive results, in which a measured physiologic response is attributed to a particular photoreceptor signal (e.g., the cones), but is in fact due to inadvertent stimulation of a different photoreceptor (e.g., the rods).

We characterized the effect of measured imprecision in the spectral output of our stimulus device (Table S2). The effect of deviations here are quite small. For example, inadvertent contrast upon the rods was <1% for the stimulus that attempted to produce isolated L+S stimulation. There is also inevitable deviation in the spectral sensitivity of the biological system from that assumed from tabular values. While estimates of population variation in opsin sensitivity and lens transmittance are available for humans, we do not have this information for the dog. In the current measurements, for example, it is possible that a large deviation from assumed biological values resulted in luminance contrast in our nominally isoluminant L–S stimulus. Future studies could vary the contrast and temporal frequency of stimulation to test for the different signatures of the chromatic and luminance pathways, and thus explore the quality of photoreceptor isolation.

We have positive evidence from our study that any inadvertent signaling from the rods was minimal, as might be expected given the photopic background light level we used^43^. In the pupil, the similar phase of response to the light flux and L+S modulations suggests that only cone signals were evoked by the light flux stimulus, and there was minimal pupil response to rod-directed stimulation. While our rod-directed stimulus also places contrast on canine melanopsin, the modulation frequency was too fast to expect a large pupil response from this mechanism^16^. In the fMRI data, the similar amplitude of response to light flux stimulation in the WT and RCD1 animals, and the absence of responses in the CRD2 model which has extensive cone degeneration at the ages studied, argues against a substantial rod contribution.

### The effects of anesthesia on pupillary and cortical responses

We collected data under general anesthesia using 100% oxygen by ventilation. The presence and level of anesthesia can influence measurements of the pupillary light reflex^49,50^. Anesthesia can also attenuate cortical responses to sensory stimulation, lowering the sensitivity of the measurement. Further, anesthesia and non-physiologic blood gasses can alter the relationships between sensory stimulation, neural response, and neurovascular coupling^51,52^. These mechanisms likely explain the paradoxical (negative) BOLD response to stimulation in the LGN we find here (and in our prior study with dogs^11^), and the difference in the sign of response between cortical and thalamic sites. While the absolute amplitude of fMRI response is readily interpretable in our data, we would ideally have a full understanding of this signal transformation.

### CRD2 is a promising model in which to study recovery of cone function

Residual responses from spared photoreceptors as seen in the RCD1 and XLPRA2 dogs (Table S1) complicates the assessment of efficacy of vision restoration treatments. For example, a small amount of intact cone function is sufficient for animals to demonstrate visual behavior and (e.g.) avoid obstacles^53^. For both behavioral and neural measures, it is easier to detect a small therapeutic effect against a consistent background of no function than it is to measure a change in a non-zero response.

Both XLPRA2 and CRD2 are associated with cone dysfunction and loss but differ in severity and tempo; in our data both show a reduction in neural response to cone-directed stimulation. We find, however, that *NPHP5*-CRD2 animals consistently show no measurable thalamic or cortical response to even the high-contrast light flux modulation. This is in agreement with electrophysiological and behavioral studies that show an absence of both rod and cone-mediated function at similar ages (Table S1). Against this background, the effect of treatment is readily apparent in the fMRI data from an individual animal, with a restoration of a normal extent of cortical response to cone-directed stimulation.

Our measure of the amplitude of pupil response was highly variable across individual animals, perhaps related to differences in pupil size and thus retinal irradiance^35^. Generally, the effect of treatment in the pupil response is measured for full-field, high-intensity flashes of light presented against a dark background^11,54,55^. Pupil responses for modulations of cone-directed stimulation around a photopic background are inevitably smaller^35^. We found in our data that fMRI measures provided more consistent results within individual animals and was more sensitive to group differences due to retinal disease.

## Author Contributions

Conceptualization: Huseyin O. Taskin, Gustavo D. Aguirre, William A. Beltran, and Geoffrey K. Aguirre; Data curation: Huseyin O. Taskin and Geoffrey K. Aguirre; Formal analysis: Huseyin O. Taskin and Geoffrey K. Aguirre; Funding acquisition: Gustavo D. Aguirre, William A. Beltran, and Geoffrey K. Aguirre; Investigation: Jacqueline Wivel; Resources: Gustavo D. Aguirre and William A. Beltran; Software: Huseyin O. Taskin and Geoffrey K. Aguirre; Supervision: William A. Beltran and Geoffrey K. Aguirre; Visualization: Huseyin O. Taskin and Geoffrey K. Aguirre; Writing - original draft: Huseyin O. Taskin, William A. Beltran, and Geoffrey K. Aguirre; Writing - review & editing: Huseyin O. Taskin, Jacqueline Wivel, Gustavo D. Aguirre, William A. Beltran, and Geoffrey K. Aguirre.

---

**Figure S1:**
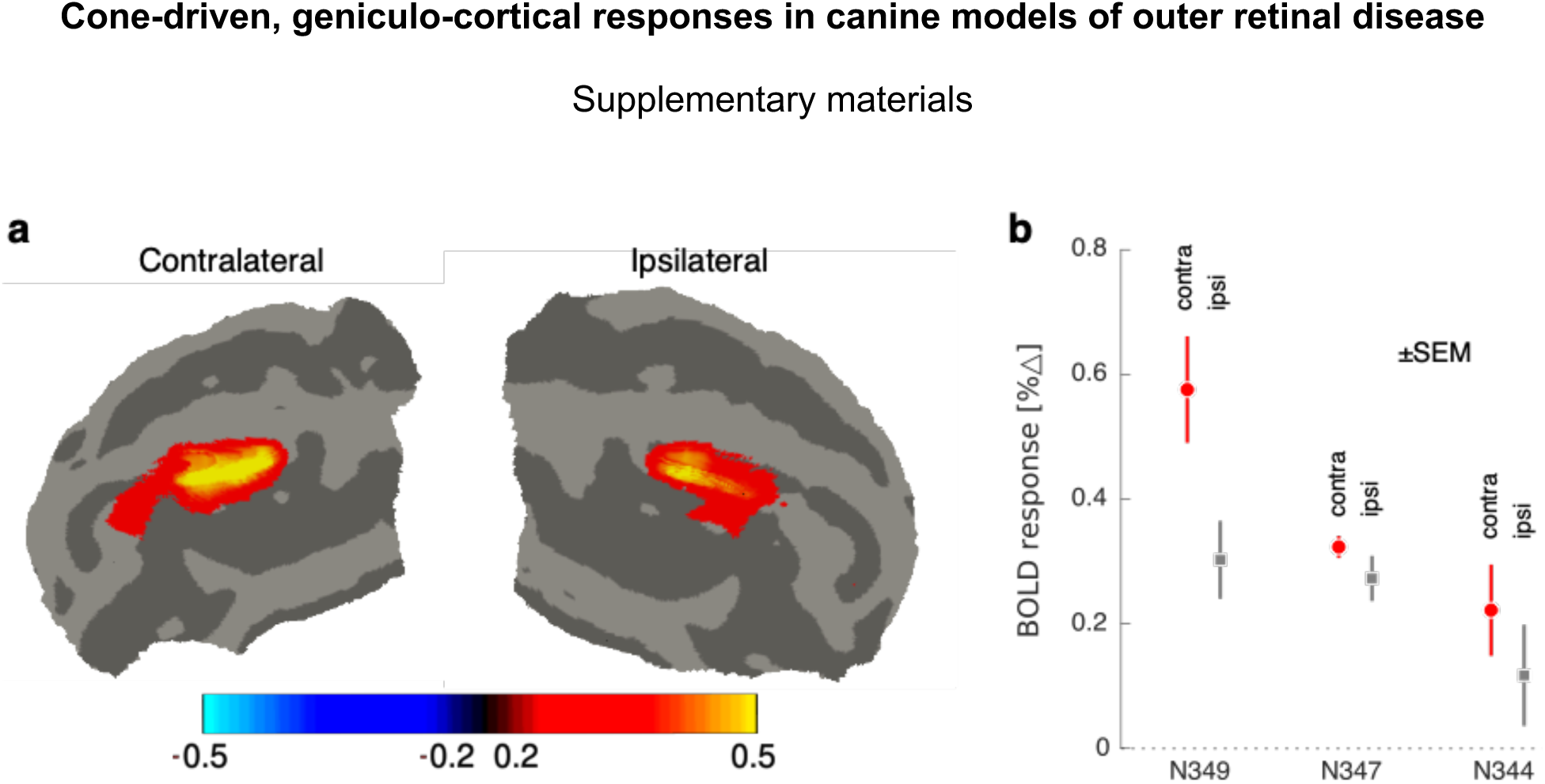
Visual cortex responses contralateral and ipsilateral to the stimulated eye. Data taken from WT animals and the light-flux stimulus. a) Surface maps showing the mean (across animals) percentage signal change to stimulation from the contralateral and ipsilateral eye. The amplitude and extent of response is slightly greater for contralateral stimulation. b) The response for each animal shows a consistent difference between the two eyes. Error bars are the standard error across acquisitions.

**Figure S2:**
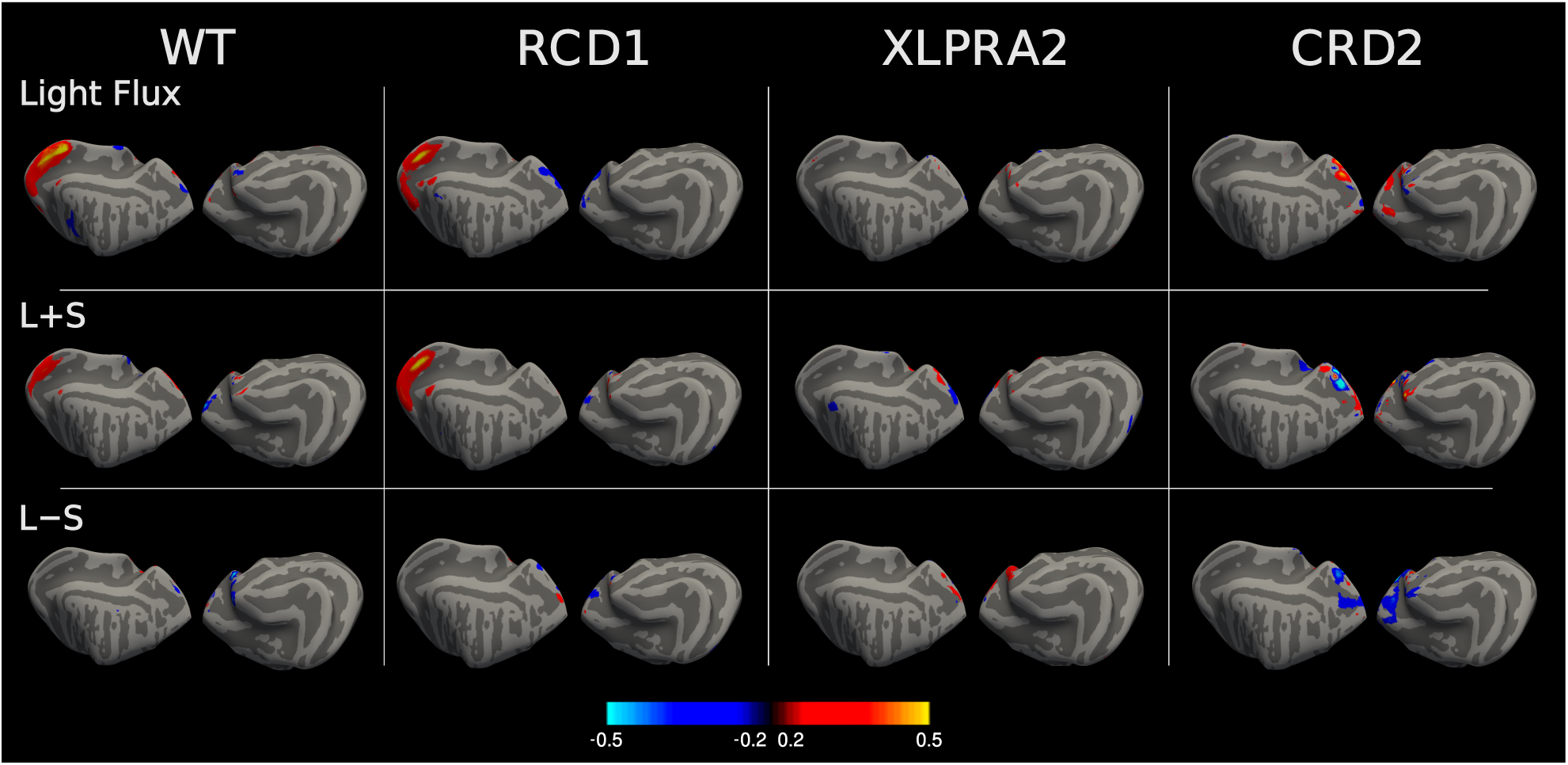
Whole brain maps of fMRI response; corresponds to Figure 5.

**Figure S3:**
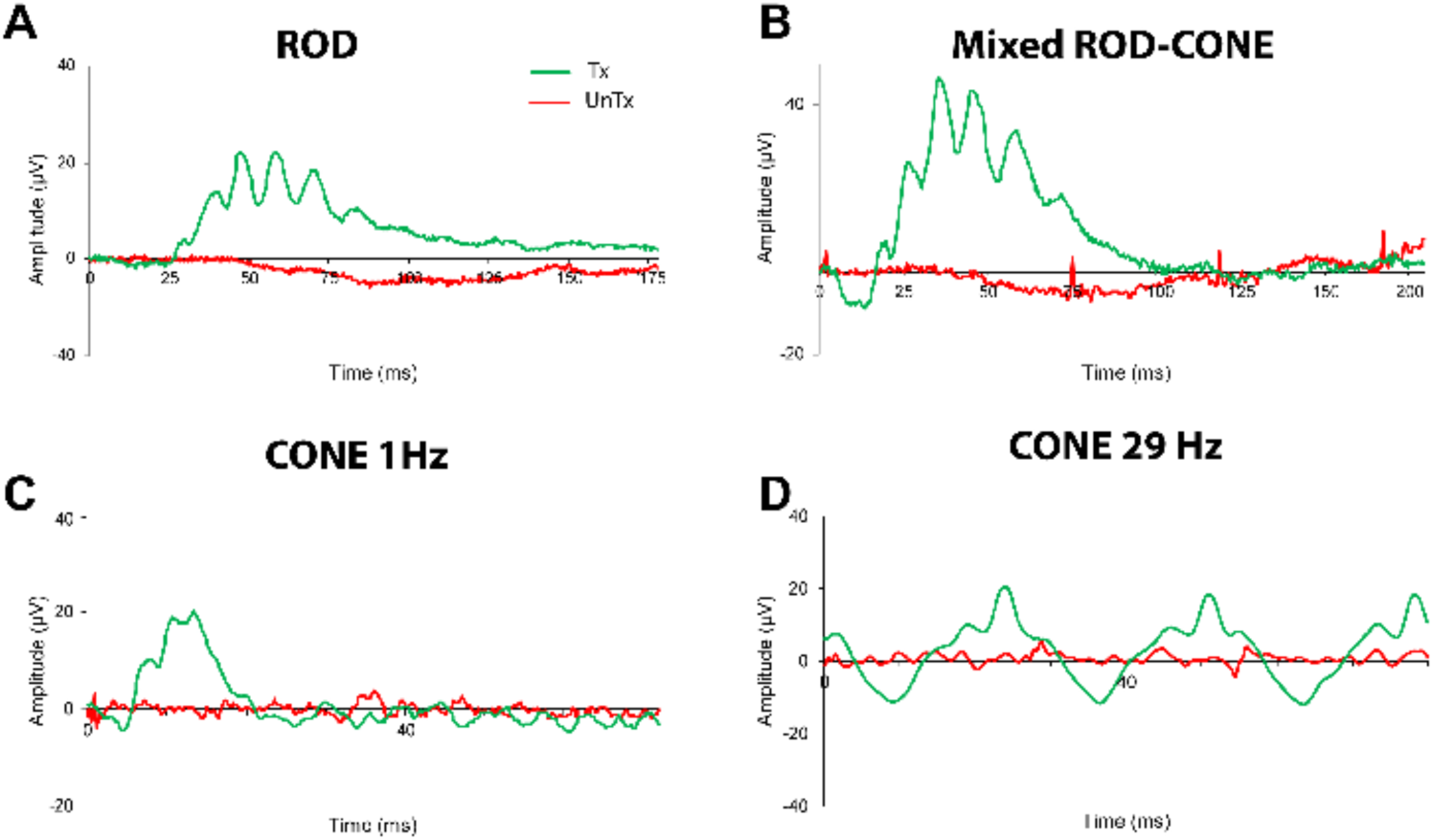
Electroretinographic recordings in a CRD2/*NPHP5* affected dog (WM67; age 38 weeks) at 24 weeks post-delivery of AAV2/5-NPHP5 to the treated (Tx, green traces) left eye. The contralateral right eye was untreated (UnTx, red traces)

**Figure S4:**
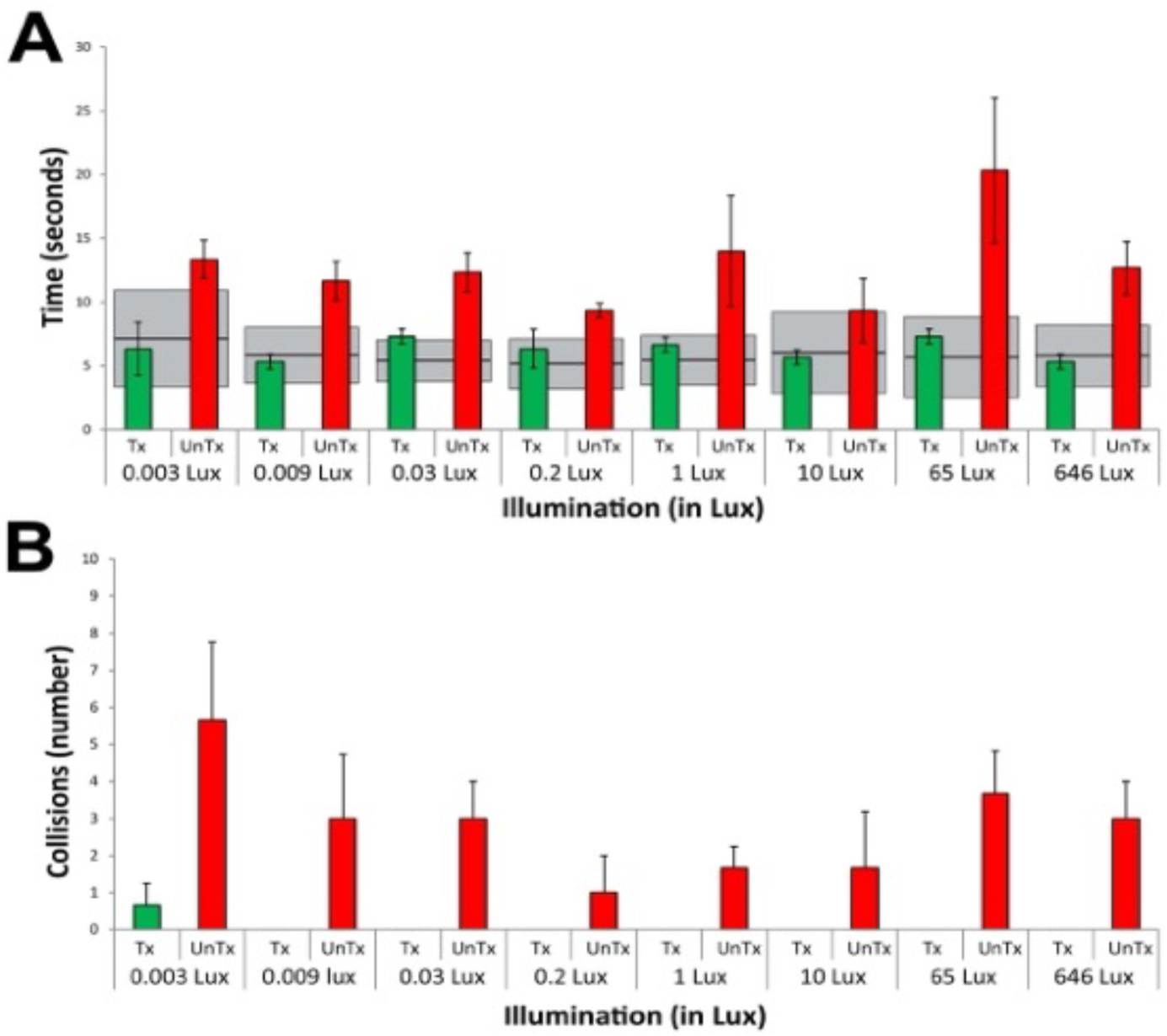
Visual function in an obstacle avoidance course under different ambient light intensities of a CRD2/*NPHP5* affected dog (WM67; age 38 weeks) at 24 weeks post-delivery of AAV2/5-NPHP5 to the treated (Tx, green bars) left eye. The contralateral right eye was untreated (UnTx, red bars) (**A**) Mean (±SD) transit time. Gray bars show 95% CI of normal (untreated) dogs. (**B**) mean (±SD) number of collisions (WT dogs have zero collisions).

**Table S1:**
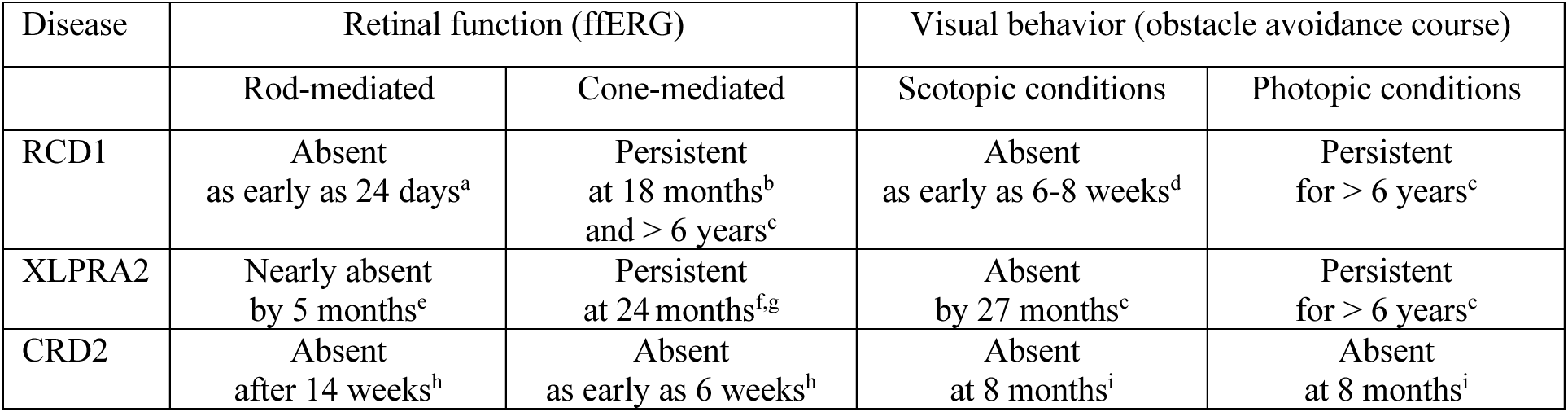
Summary of ages at which retinal and visual function are detectable in RCD1/*PDE6B*, XLPRA2/*RPGR*, and CRD2/*NPHP5* affected dogs. ffERG: full field ERG ^a^: Aguirre GD, Rubin LF. Rod-cone dysplasia (progressive retinal atrophy) in Irish setters. *Journal American Veterinary Association*, 1975; 166 (2): 157-164. ^b^: Petit L, Lheriteau E, Weber M, et al. Restoration of vision in the pde6beta-deficient dog, a large animal model of rod-cone dystrophy. *Molecular Therapy*, 2012: 20 (11): 2019-2030 ^c^: Beltran WA (personal communication) ^d^: Hodgman SFJ, Parry HB, Rasbridge WJ, Steel JD. Progressive Retinal Atrophy in dogs 1. The disease in Irish Setters (Red). *Veterinary Record*, 1949; 61 (15): 185-189. ^e^: Dufour VL, Cideciyan AV, Ye G-J, et al. Toxicity and efficacy evaluation of an adeno-associated virus vector expressing codon-optimized RPGR delivered by subretinal injection in a canine model of X-linked Retinitis Pigmentosa. *Human Gene Therapy*, 2020: 31 (3-4): 253-267. ^f^: Beltran WA, Cideciyan AV, Iwabe S. Successful arrest of photoreceptor and vision loss expands the therapeutic window of retinal gene therapy to later stages of disease. *Proceedings of the National Academy of Sciences of the USA*, 112 (43): E5844-5853. ^g^: Beltran WA, Cideciyan AV, Boye SE, et al Optimization of retinal gene therapy for X-linked retinitis pigmentosa due to RPGR mutations. *Molecular Therapy*, 2017; 25 (8): 1866-1880 ^h^: Downs LM, Scott EM, Cideciyan AV et al. Overlap of abnormal photoreceptor development and progressive degeneration in Leber congenital amaurosis caused by NPHP5 mutation. Human Molecular Genetics, 2016; 25 (19): 4211-4226 ^i^: Aguirre GD, Cideciyan AV, Dufour VL et al. Gene therapy reforms photoreceptor structure and restores vision in NPHP5-associated Leber congenital amaurosis. *Molecular Therapy*, 2021; 29 (12): 2456-2468

**Table S2:**
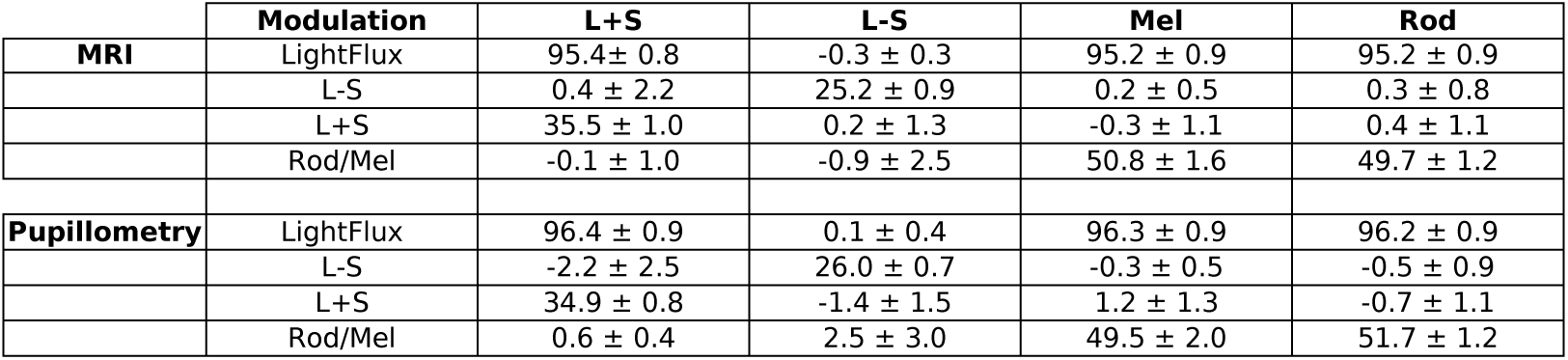
Spectroradiometric measurements of the stimuli were made after each experimental session. Inevitable imprecision in device control leads to variation in contrast on the targeted photoreceptor populations. Shown is the mean and standard deviation across animals of the calculated contrast upon the targeted and silenced photoreceptor classes for the MRI and pupillometry studies.

